# Quantifying Intracellular Mechanosensitive Response upon Spatially Defined Mechano-Chemical Triggering

**DOI:** 10.1101/2025.05.04.652118

**Authors:** Elaheh Zare-Eelanjegh, Renard T.M. Lewis, Ines Lüchtefeld, Ulrike Kutay, Tomaso Zambelli

**Affiliations:** Laboratory of Biosensors and Bioelectronics, Institute for Biomedical Engineering, ETH Zurich, Zurich, Switzerland; Department of Biology, Institute of Biochemistry, ETH Zurich, Zurich, Switzerland

**Keywords:** Mechanotransmission, nuclear mechanoresponse, endoplasmic reticulum mechanoresponse, tension, lamins, cytoskeleton

## Abstract

The mechanotransduction process relies on the interaction of mechanical and biochemical cues, transmitting cellular forces to intracellular organelles to activate biochemical pathways and elicit responses. This involves mechanoresponsive components like actin filaments, microtubules (MTs), and the lamin meshwork. Fluidic force microscopy (FluidFM), a force-controlled micropipette allows for the manipulation of intact cells mechanically and chemically, providing a novel approach to study mechanotransmission in cells *in situ*. FluidFM combined with fluorescence lifetime imaging microscopy (FLIM), enables high-resolution mapping of intracellular tension dynamics. Here, we used cells with varying nuclear lamina compositions to explore the lamina’s role in initiating mechanoresponse to external cues. We found that A-type and B-type lamins trigger nuclear mechanoresponse distinctly, with A-type lamins contributing to nuclear elasticity, whereas B-type lamins influence viscous response. Moreover, MTs underwent mechanical adaptation and assisted in releasing the tension in lamin A/C knockout (KO) cells, contrasting with healthy cells where MTs aid to preserve the tension locally rather than transferring it. This research provides insights into the dynamic mechanoresponse of cellular components and supports targeted therapies for mechanical stress-related diseases.

## Introduction

Mechanical forces regulate cell behavior and function. The cytoskeleton, particularly the actinomyosin network, acts as a primary receptor and transmitter of these signals.^1,2^ Biological tissues and organs are influenced by these mechanical signals at the cellular level.^3^

Moreover, cells coordinately transmit external forces into the intracellular organelles, activate biochemical pathways, and eventually elicit corresponding cellular responses.^4,5^ This relies on interconnected mechanosensory systems, including ion channels, membrane receptors, cytoskeletal components, and the Linker of Nucleoskeleton and Cytoskeleton (LINC) complex that bridges the cytoskeleton to the nucleus.^6^ Additionally, the nucleus, a major mechanosensitive organelle, can directly sense large-scale forces resulting in nuclear deformations and gene expression changes via a process called ‘nuclear mechanotransduction’.^7^ Other organelles such as the endoplasmic reticulum (ER)^8^, mitochondria^9^ and Golgi apparatus^10^ can also sense transmitted forces. The nuclear lamins, intermediate filaments lining the inner side of inner nuclear membrane, likely collaborate with the LINC to regulate nuclear tension and mechanotransduction.^11,12^ Forces convert into biochemical cues at the nuclear envelope through phosphorylation of lamin A and emerin, a mechanosensitive inner nuclear membrane (INM) protein that associates with lamin A/C.^13^ The general belief is that viscoelastic responses of the nucleus to physiological forces are governed by the interactions between A- and B-type lamins as well as chromatin.^14^ Understanding the mechanical roles of lamin A/C and lamins B1/B2 will enhance our knowledge of nuclear deformation in health and disease. Notably, different mechanotransduction pathways crosstalk and interplay with biochemical signals (*e.g.*, growth factors) to elucidate specific cell behaviors.^15^ Disruption of biomechanical homeostasis and failure of cells to appropriately transmit the mechanical stimuli into the requisite biochemical pathways, can lead to diseases such as arthritis, atherosclerosis, and cancer.^16,17^

Several methods have been developed to explore single cell mechanics and mechanotransmission including atomic force microscope (AFM), optical tweezer (OT), magnetic tweezer (MT), biomembrane force probes (BFP), traction force microscope (TFM), optogenetic mechanostimulators and DNA-based mechanical sensors.^18,19^ Recently, a time-shared optical tweezer microrheology method (TimSOM) that measures viscoelastic properties of cytoplasm, nuclear envelope, and nucleoplasm in living systems has been reported.^20^ However, these methods often face limitations in measuring intracellular forces at the molecular level within the cell’s natural state.^21^ For instance, the recently developed TimSOM requires embedding tracer beads inside cells or embryos and yields averaged measurements over small regions rather than pinpointing single molecular-scale forces. To overcome such constrains, FRET^22,23^ and FLIM-based^24,25^ tension sensors have been engineered for studying intracellular mechanosensing at the molecular level. FRET detects energy transfer between two fluorophores in proximity, indicating molecular interactions and conformational changes within biological systems. Unlike intensity-based FRET methods, FLIM measures the lifetime of fluorescent molecules excited by a laser pulse, offering a quantitative signal for molecular interactions independent of reporter molecule concentration. To delve into the interplay of biochemical and mechanical aspects in cellular behaviors, we require tools that can sensitively stimulate living cells both mechanically and chemically in their natural state, while observing intracellular dynamics. Furthermore, these methods should quantify minute forces with superior spatial and temporal resolution.

Fluidic force microscopy (FluidFM), a micropipette configured on an AFM, addresses this challenge.^26^ The FluidFM micro channeled-cantilever with a nano-aperture, connected to a fluidic pressure controller, enables quantitative injection of impermeable molecules into subcellular compartments of living cells.^27-30^ such injections can be realized as biochemical triggering of the cellular events with spatial and temporal resolution while preserving cell viability. Combining FluidFM with optical readouts like FLIM-based molecular sensor enables simultaneous manipulation of intact cells and quantifying their responses.

In this work, we first used FluidFM combined with the FLIM for *in situ* mechanical (*i.e.*, indentation) and chemical (*i.e.*, drug injection) triggering of cells with simultaneous intracellular tension mapping at the nuclear envelope and ER. Further, mechanoresponsive crosstalk between different organelles within the cell including the nucleus, ER and cytoskeleton. For chemical interference, we injected cytochalasin D (CytoD) and/or nocodazole (Noco) to inhibit the actin and/or microtubule network, respectively. Next, we exploited cell lines in which nuclear lamins were perturbed (*i.e.*, wild-type HeLa (WT), lamin A/C gene (*LMNA)* KO, *LMNB1* and *LMNB2* (LMNB)-knockdown (KD) and *LMNA* KO+LMNB-KD and subjected to mechano-chemical stimulation and quantification of tension changes.

## Results and discussion

We had previously reported the development of a FluidFM-FLIM system based on a customized stage for mounting our AFM head on top of a confocal inverted microscope to investigate the effect of the plasma membrane tension on the opening of mechanosensitive ion channels.^31^ Here, we used HeLa cells stained with ER Flipper-TR, a tension reporter dye which selectively labels the ER and nuclear membrane constituting a continuous membrane network.^24^ ER Flipper-TR measures the membrane tension changes through alterations in its fluorescence lifetime and serves as a quantitative readout for the tension responses to the external stimuli within living cells. FLIM signals were recorded before, during and after the mechanical or mechano-chemical stimulus facilitated by FluidFM.

### Mechanical stimulation of the nucleus: impact of probe apex shape on cellular mechanoresponse

The geometry of an AFM probe determines how a force is spatially applied on a cell surface.^32^ Different probe shapes cause varying membrane deformations, activating diverse mechanoresponses through mechanical involvement of various organelles. The cellular cytoskeleton plays crucial roles in distributing and absorbing mechanical forces to maintain cellular integrity and function. Cylindrical and pyramidal FluidFM probes were used for indenting and puncturing the nucleus under varied forces (Fig. 1 A and B). Cylindrical FluidFM probes were sharpened using a focused ion beam (FIB) to facilitate membrane rupture with minimal stress (Fig. 1B, see Methods section). In parallel, Lifetime images of ER Flipper-TR stained HeLa cells were obtained using a FLIM detector integrated into a confocal laser scanning microscope (Fig. 1C). The images, color-coded for clarity, report the conformational shifts in the ER Flipper-TR molecule. Higher lifetime values (red color) indicate increased membrane tension.

**Fig. 1.**
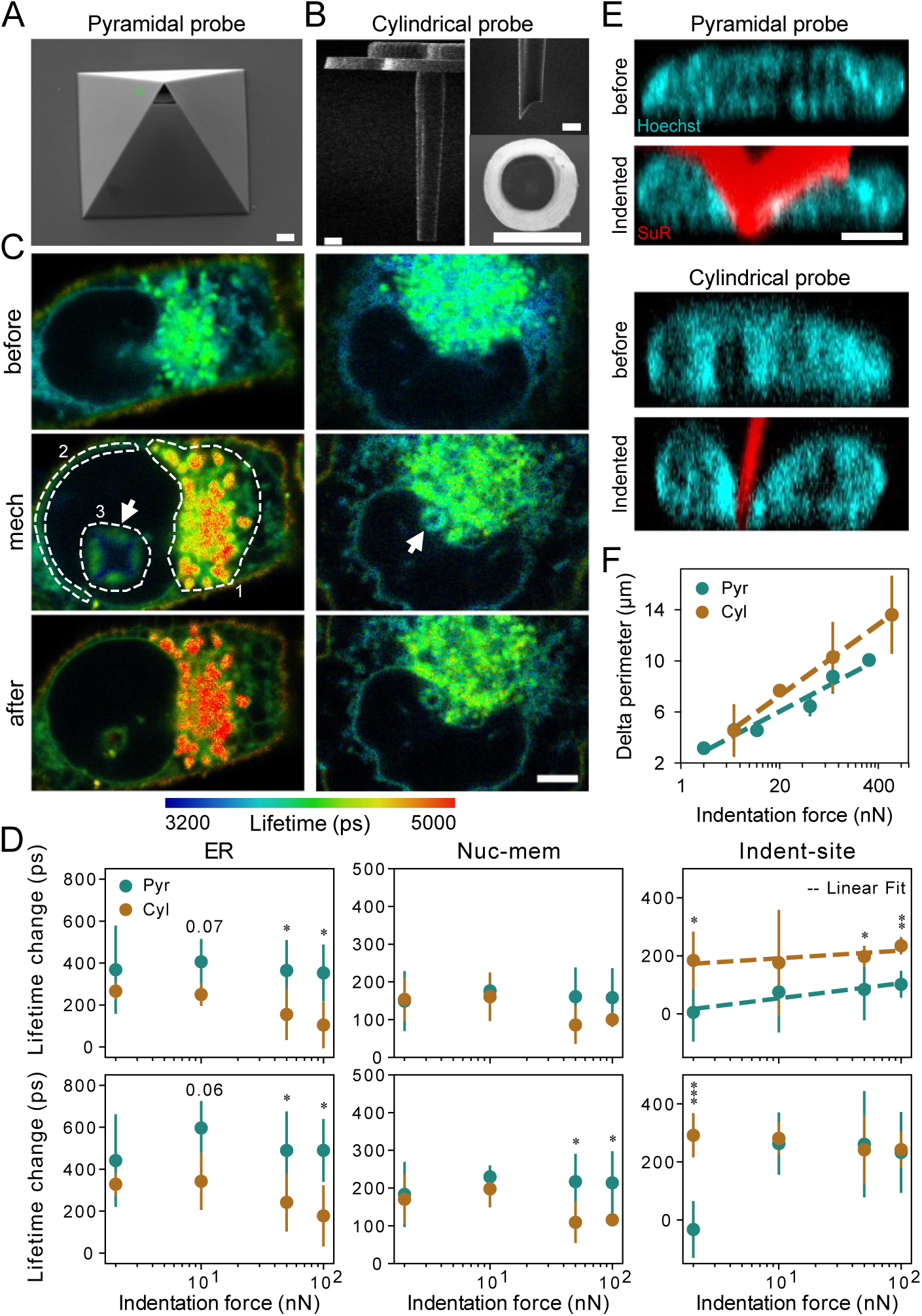
FluidFM stimulation combined with FLIM readout for measuring subcellular mechanoresponse upon mechanical stimulation of nuclei. **A, B)** SEM micrographs of a pyramidal probe (A) and a cylindrical probe (B) before (left) and after FIB-cut (top right). The top view of the cylinder is also shown in panel B. Scale bars represent 1 µm. **C)** Lifetime images of HeLa cells stained with molecular tension probe (ER Flipper-TR) before (top), during (middle) and after (bottom) indentation at 100 nN, with pyramidal (left panel) and sharpened cylindrical probes (right panels). Arrows denote the indentation site when either pyramidal or cylindrical probes are used. Dashed lines indicate a representative selection of ROIs, encompassing 1) the ER, 2) nuclear membrane (Nuc-mem), and 3) the area around the indentation site (Indent-site). Scale bar represents 5 µm. **D)** Change in lifetime during (top panel, “mech” in C) and after (bottom panel) mechanical stimulus with varied indentation forces at different ROIs: ER, nucleus membrane (Nuc-mem) and incident indentation site (“Indent-site”) (Mean ± SD, N ≥ 2, n ≥ 4, ***p < 0.001, **p < 0.01, *p < 0.05, reported p-values of 0.05 ≤ p < 0.1, t-test). For mechanical stimulation, pyramidal (“Pyr”) and cylindrical (“Cyl”) probes with similar stiffness (2 ± 0.5 Nm-1) were utilized. **E)** Lateral view of nucleus before (top panels) and during indentation (bottom panels) (indicated by “mech” in panel C) at 100 nN using pyramidal and cylindrical probes. Nuclei are stained with Hoechst dye (blue) and probes are filled with sulforhodamine dye (SuR, red). Scale bar represents 5 µm. **F)** Change in perimeter of nucleus measured from confocal images at varied indentation forces, with pyramidal and sharpened cylindrical probes (Mean ± SD, N = 2, n ≥ 3).

Subsequently, different “Region of Interests” (ROIs) were manually selected for the ER, nuclear membrane (Nuc-mem), and the area around the indentation site (Indent-site) (*SI Appendix*, Fig. S1 and Fig. 1C middle) and their lifetime values were analyzed. Mean lifetime values within each ROI were normalized by subtracting the initial values for each cell (see *SI Appendix* and Fig 1D). The average lifetime values at the ER showed greater tension increase with pyramidal probes compared to cylindrical probes across a range of indentation forces (10 - 100 nN) (Fig. 1D, top left). The ER retains this higher tension even after retracting the probe and removal of the external mechanical stimulus (Fig. 1D, bottom left). At the nuclear envelope (NE), both pyramids and cylinders impose the same amount of stress during mechanical stimulus (Fig. 1D, top middle). Yet, following the removal of the probe terminating the mechanical perturbation, pyramidal probes reveal increased tension at NE at higher indentation forces of 50 nN and 100 nN in contrast to the cylinders (Fig. 1D, top middle). At the NE, a decrease in lifetime or relaxation in tension occurred between 10 nN and 50 nN indentation, observed only with cylindrical probes. Since this is within the expected range of indentation force during membrane rupture, the observed relaxation in tension is related to membrane rupture and cylinder insertion in HeLa cells. Although pyramidal probes are also expected to puncture the cell membrane and subsequently the NE upon indenting cells targeting their nucleus with increasing forces (*i.e.*, around 100 nN for HeLa cells^28^), their insertion phenomena within the membranes is not instantaneous and thus not detectable within the graph demonstrating the quantified tension upon indentation (Fig. 1D, top middle). Sharpened cylindrical probes elicit higher tension locally at the indentation site when compared to pyramids during indentation (Fig. 1D, top right). This is due to the cylindrical geometry leading to a smaller area where the inserting force is employed, increasing the pressure locally, while minimizing the disturbance to other organelles such as the ER. Hence, sharpened cylindrical probes exert a lower overall tension on the cells during injection. Immediately following the retraction of cylindrical probes, the tension remains constant or slightly reduces. But, for pyramidal probes, surprisingly, the lifetime increases within all ROIs immediately following the retraction. This suggests an additional mechanical stress upon the withdrawal of pyramidal probes.

Lastly, to assess the origin of higher tension imposed locally by cylindrical probes compared to pyramidal ones, the deformation of cell nuclei was quantified during the indentation process using lateral images acquired by the confocal microscopy (Fig. 1E). For this, FluidFM probes were filled with sulforhodamine (SuR) dye (red) and the nuclei, stained with Hoechst (blue) were subjected to indentation at varied forces while acquiring z-stacks images before and during stimulus. Then, the lateral views were obtained from z-stack images, and the nucleus perimeter was measured before and during indentation for both pyramidal and cylindrical probes (Fig. 1F). As expected, cylinders introduce higher deformation of cell nuclei with maximum deformation around the apex (Fig. 1E). This rationalizes the higher lifetime values reported for the indentation site for cylinders compared to pyramids (Fig. 1D, top right).

Collectively, the use of different probe shapes for mechanical stimulation of the nucleus revealed a distinct tension distribution across varied subcellular compartments. While tension was higher around the cylindrical probe, a higher tension at the ER was observed when pushing with a pyramidal probe. This could be attributed to the greater total volume displacement caused by the broader apex of pyramidal probes compared to the sharper tips of cylindrical ones. Furthermore, tension was increased in the ER and the NE of cells stimulated by the pyramids upon removal of probes, which was not the case for the cylinders. Since the cytoskeleton is essential for sensing such mechanical stimuli and for the downstream responses (*e.g.*, transcription), we further explored whether chemically induced depolymerization of actin filaments affects the response of cells to mechano-stimulation, combining nucleus indentation and CytoD injection.

### Disrupting actin filament dynamics by mechano-chemical triggers inhibits their active support of the stressed nucleus

The dynamics of cytoskeletal and nuclear actin, whether in monomeric (G-actin) or filamentous (F-actin) forms, affects both the sensing and response to mechanical stimuli, as well as the resulting transcriptional processes.^33,34^ The mechanosensitive LINC complex in tandem with cytoskeletal components regulates nuclear function and protects the nucleus from excessive mechanical stress. CytoD disturbs actin filament dynamic mainly by binding to G-actin and inhibiting filament assembly.^35^ We investigated whether CytoD interruption of the actin network alters the tension load on the nucleus, thus modifying the nuclear mechanoresponse. Therefore, actin filaments were chemically disrupted by a mechano-chemical stimulation through indenting the nucleus and concurrently injecting CytoD in a spatio-temporally defined fashion (Fig. 2 A and B). Since CytoD is smaller in size than the passive diffusion limit of nuclear pore complexes, it distributes throughout the nucleus and cytosol, interacting with both nuclear and cytoskeletal actin, thereby influencing overall cell structure and its mechanoresponsive behavior. While CytoD is believed to permeate cell membranes, its active intracellular injection allows for real-time tracking of dynamic responses. Additionally, when quantitatively injected, the amount taken up by different cells is consistently controlled, regardless of variations in cell responses.

**Fig. 2.**
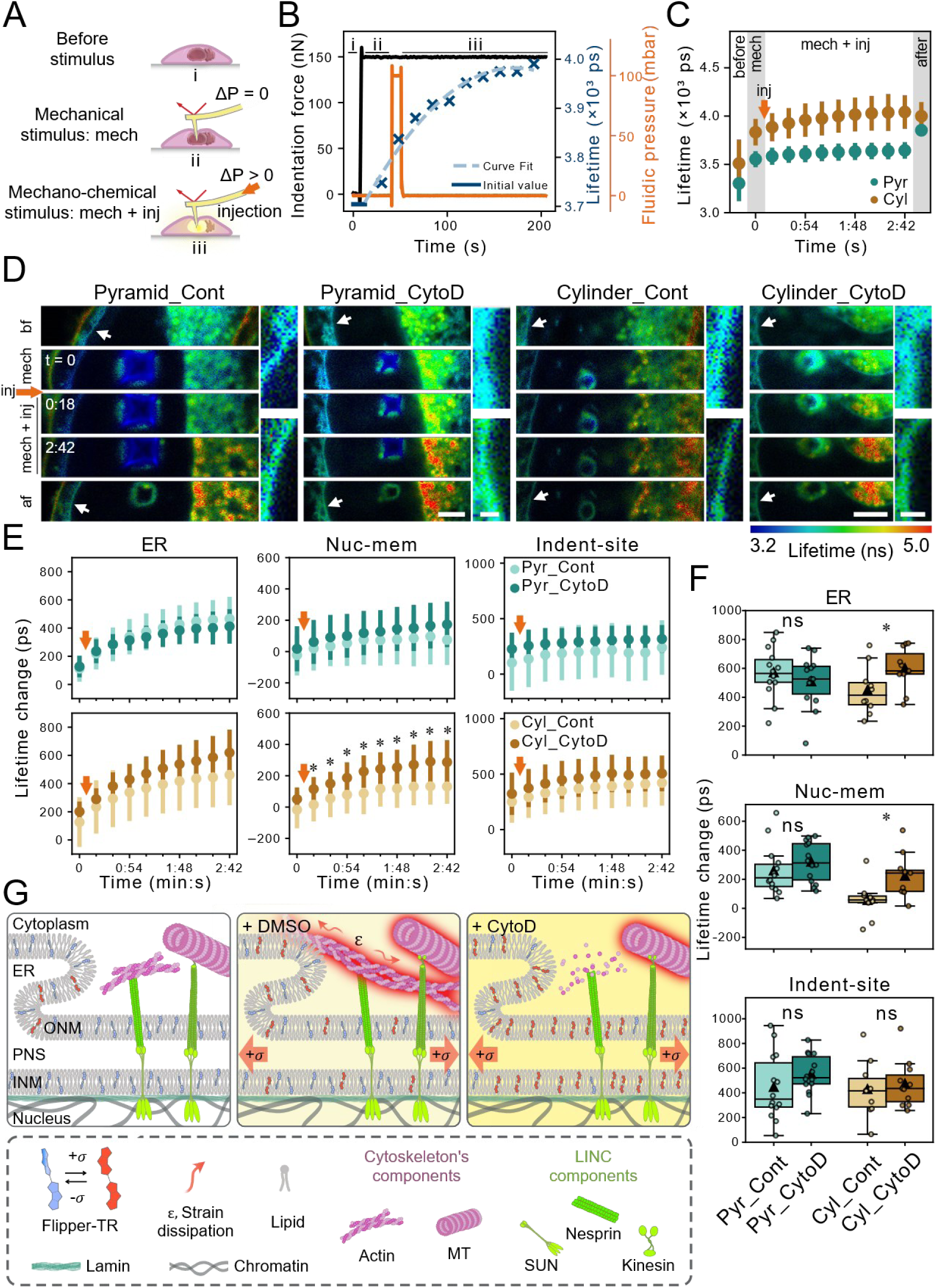
FluidFM-FLIM mechanochemical stimulation of HeLa cell nuclei using pyramidal and cylindrical probes. **A)** Schematic showing the mechanotransduction at LINC complexes before (top) and after external stimulus: mechanical (middle) or mechanochemical (bottom) stimulus. ΔP represents the fluidic pressure difference between the probe reservoir and the aperture. When this pressure difference is positive, it indicates that the injection mode of FluidFM is being utilized. **B)** Concurrent readout of AFM-measured indentation force, ER Flipper-TR lifetime, and fluidic pressure pulse for injection over time. For assured injection, 150 nN was chosen as the set point for indentation using both probe geometries. Parts i, ii and iii on the graph in B and the schematic in A represent: i) before stimulus, ii) during mechanical stimulus when the probe has reached the set point on the cell without fluidic pressure application, and iii) during mechanochemical stimulus when the probe remains in contact with the cell at the set point, and a fluidic pressure pulse is applied for drug (or control) injection. **C)** Absolute lifetime values before, during and after injecting HeLa cells with CytoD with pyramidal (“Pyr”) and cylindrical (“Cyl”) probes at indentation site (Mean ± SD, N ≥ 3, n ≥ 11). The orange arrow indicates the injection pulse at the specified time in the mechanochemical triggers. **D)** Time-lapse lifetime images of ER Flipper-TR-stained HeLa cells before, upon indentation (“mech”), after indentation plus injection (“mech + inj”) of either DMSO (as control) or 50 µM CytoD, and after retracting pyramidal (left) and cylindrical (right) probes. The numbers represent time (min:s), with time 0 corresponding to the indented state (mechanical stimulus, noted as “mech”). Inserts represent magnified nuclear membrane at locations specified with arrows. Scale bars in time-lapse images and inserts represent 5 µm and 1 µm, respectively. **E)** Change in lifetime at different ROIs: ER, nucleus membrane (Nuc-mem) and indentation site (“Indent- site”) when mechanochemically challenged by either CytoD or Control using pyramidal (top row) and cylindrical (bottom row) probes (Mean ± SD, N ≥ 3, n ≥ 11, ***p < 0.001, **p < 0.01, *p < 0.05, reported p- values of 0.05 ≤ p < 0.1, t-test). Orange arrows indicate the injection pulse at the specified time in the mechanochemical process. **F)** Box plots showing lifetime change upon retracting probes from the injected cells at different ROIs: ER, Nuc-mem and indentation site. Mean values are shown with triangles (N ≥ 3, n ≥ 11, ***p < 0.001, **p < 0.01, *p < 0.05, reported p-values of 0.05 ≤ p < 0.1, ns: not significant, t-test). **G)** Schematic represents the tension report of ER Flipper-TR dye inside the membrane of the nucleus and the ER before (left panel) and after mechanochemical triggering of the nucleus with DMSO (as control, middle panel) or CytoD (right panel). In contrast to control cells, delivery of CytoD induces more accumulated tension (i.e., FLIM data) at the nuclear membrane and the ER because it disrupts load-bearing actin component of cytoskeleton. ONM: outer nuclear membrane, INM: inner nuclear membrane, PNS: perinuclear space, and σ: tension exerted by indenting the nucleus using FluidFM probes.

FluidFM probes were filled with 50 µM CytoD in DMSO (or DMSO for control groups) and used for mechano-chemical triggering by indenting the nucleus at 150 nN (“mech”), followed by a quantitative intracellular injection of CytoD or DMSO (“mech + inj”) (Fig. 2 A and B). The indentation force was adjusted to 150 nN to ensure a successful injection into HeLa cells. FLIM measurements were performed before, during “mech” and “mech + inj”, and just after the manipulation (Fig. 2 B and C). Indentation force applied by AFM, serving as a controlled mechanical stimulus (black line), fluidic pressure pulse used for injection (orange line), and time-lapse lifetime values recorded by the FLIM detector (dark blue crosses and light blue fitted curve) were recorded simultaneously (Fig. 2B). This injection process delivers ∼10 fL of CytoD (or DMSO), equivalent to ∼1% of the cell’s volume, leading to a final intracellular concentration of 0.5 µM. During “mech + inj”, the probe remained inside the cell for 3 min ± 10 s. After this period, the probe was retracted, and the lifetime values of the cell were captured again. Next, we compared responses under mechano-chemically challenging conditions: injection of CytoD (Fig. 2C) or DMSO (*SI Appendix*, Fig. S3). A markedly higher tension at the indentation site was observed for cylindrical probes compared to pyramidal ones. As observed before (Fig. 1D), cylindrical probes exert a higher tension at the indentation site both with and without CytoD injection. In the case of pyramidal probes, tension increased at the indentation site at “mech” state but remained constant for the entire “mech + inj” state. In contrast, by cylindrical probes tension increases at the indentation site during the “mech” state and increases with time after injection of CytoD (“mech + inj”). After removal of the stimulus, tension at the indentation site shows an increase for pyramidal probes and a slight decrease for cylindrical probes, consistent with the results shown in Fig 1D.

Time-lapse lifetime images of representative cells were recorded at various stages: before, during mechanical stimulus (“mech”), after mechanical stimulus and injection (“mech + inj”), and post-retraction, using pyramidal and cylindrical probes (Fig. 2D). Lifetime changes were calculated by subtracting each cell’s initial lifetime value and averaging these changes for each condition (CytoD or Cont), both during (Fig. 2E) and after the stimulus (Fig. 2F). CytoD-injected and control cells showed a similar tension across all the ROIs in the case of pyramidal probes (Fig. 2E, top row). For cylindrical probes, CytoD-injected cells exhibited a slight increase (*p*-value of 0.087) at the ER and a significant increase (*p*-value of 0.015) at the NE when compared to control cells (Fig. 2E, bottom row). Interestingly, after retracting probes (Fig. 2F), tension at the ER and the NE remained higher for CytoD-treated cells compared to control groups when challenged by cylindrical probes. By retracting pyramidal probes, similar levels of tension were observed for CytoD- and control groups within all the ROIs. In agreement with Fig. 1D, retracting pyramidal probes imposed an additional mechanical stress onto the cell (Fig 2 E and F).

Elevated tension at the NE and indentation site is likely linked to actin filaments loss in CytoD-injected cells, as actin filaments known to protect nuclei from the excessive mechanical stress.^36^ Reportedly, external mechanical stimulation of cells as well as intensive stretching of the NE during cell migration trigger a transient formation of an actin rim around the nucleus or actin cap which is hypothesized to protects nuclear content from mechanical damage by minimizing the nuclear deformation.^37^ Therefore, hindering actin polymerization by CytoD injection inhibits actin cap formation which otherwise dissipates the excessive strain thus lowering the tension as a response to the external stimuli (Fig. 2G).

Furthermore, the ER displayed an increasing tension with time. As the NE is contiguous with the ER, stretching and deforming the nucleus is likely to increase the membrane tension in the neighboring ER^38^. Additionally, the ER extends throughout the entire cell. Therefore, when targeting the nucleus even with sharpened cylindrical probes, the ER is likely affected as well. However, similar tension at the ER and NE for CytoD-injected and control cells during and after stimuli were observed when pyramidal probes were used (Fig. 2F). The pyramidal probes with a bigger apex size compared to cylindrical ones insert a mechanical disturbance directly to the ER and the NE while targeting the nucleus. This results in similarly elevated tensions for both control and CytoD-injected cells when stimulated with pyramidal probes.

Additionally, the interplay between actin dynamics and mechanotransmission at the ER and NE highlights the complexity of cellular mechanoresponses and emphasizes the need for further investigation into their cross-reactivity and directionality. Thus, we applied mechano-chemical triggering to a variety of target organelles, including the peripheral ER, nucleus, and cell periphery and further studied the tension propagation *in situ* as a response to these stimuli.

### Interconnected mechano-crosstalk between ER, nucleus, and actinomyosin: investigating the impact of target organelle for the mechano-chemical stimuli

The mechanical crosstalk between the ER, nucleus, and cytoskeleton is complex because of their intricate interconnection.^39^ While it is acknowledged that the nucleus serves as an endpoint for the dispersion of forces, there is limited research specifically addressing cross-reactivity and directionality of mechano-signaling between these subcellular components. Therefore, a cylindrical probe was used for indentation on either the peripheral ER, the nucleus (with minimal ER lumens above) or the cell periphery with limited ER involvement (Fig. 3 A and B). Besides, intracellular injection of either CytoD or DMSO (control) was performed. In parallel, the tension at the ER and the NE of cells was monitored during (Fig. 3C) and after (Fig. 3D) mechano-chemical triggering. When the nucleus was targeted (Fig. 3C, top row), CytoD-injected cells showed higher tension at both the ER and NE compared to the control group. Conversely, for cell periphery-targeted cells (Fig. 3C, bottom row), control cells showed a higher tension at both the ER and NE compared to CytoD-injected cells. Interestingly, when the ER was subjected to the same stimuli (Fig. 3C, middle row), no detectable change was observed between the cells injected with or without CytoD. Following the probe retraction, tension at the ER and NE was higher for CytoD-injected compared to control cells when the ER and nucleus were targeted (Fig. 3D). In contrast, in cells targeted for the cell periphery, slightly lower tension at the ER and similar tension at the NE were observed for CytoD-treated compared to control cells.

**Fig. 3.**
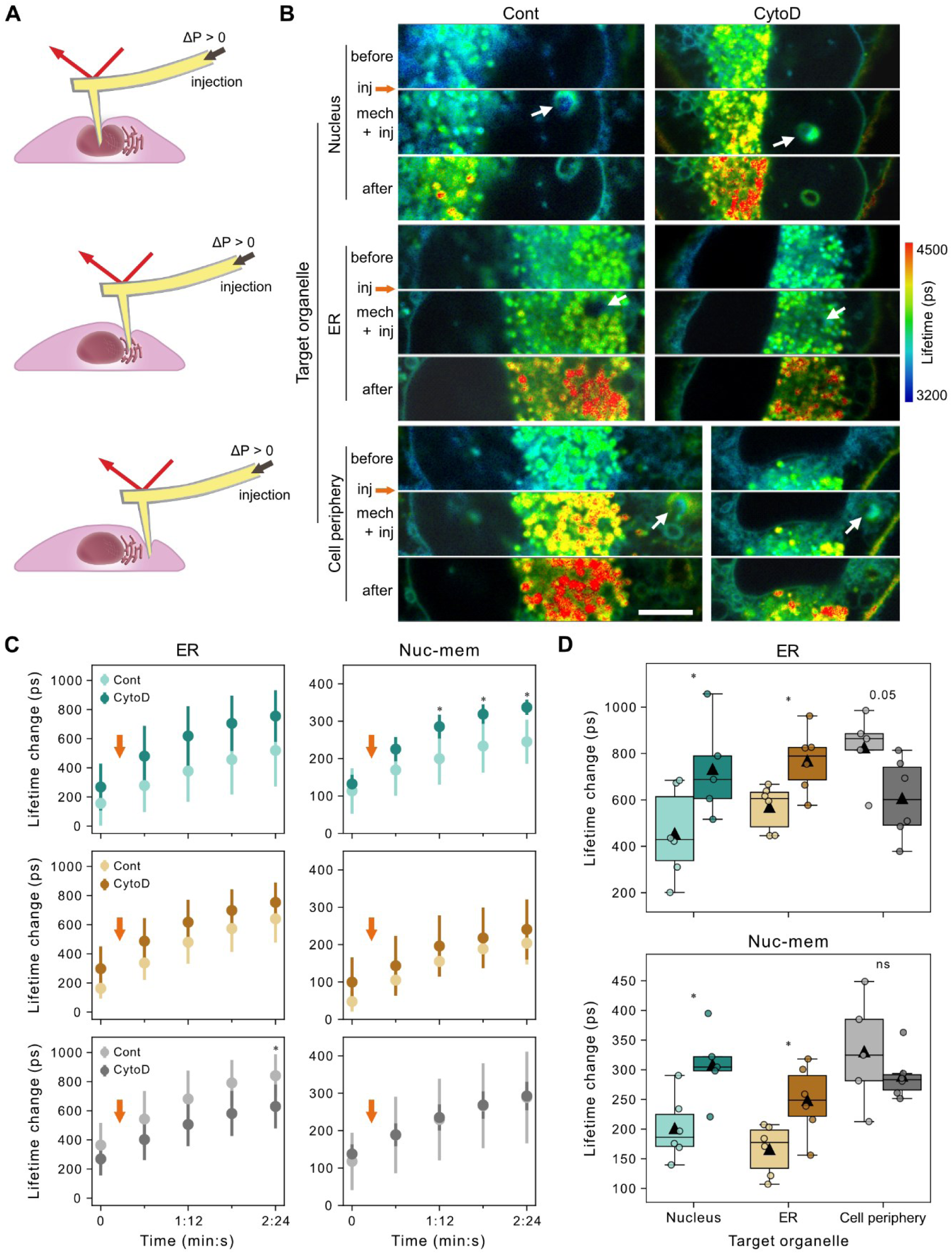
ER and nuclear envelope crosstalk as a response to mechanochemical triggering of different intracellular components of HeLa cells. **A)** Schematic shows targeting different organelles for mechanochemical stimulation: nucleus (top), the ER (middle) and cell periphery (bottom). **B)** Lifetime images of HeLa cells stained with the ER Flipper-TR before, at mechanochemical stimulated state (“mech + inj”) and after stimulation targeting different subcellular compartments: nucleus (top panel), the ER (middle panel) and cell periphery (bottom panel) for indentation and injection of either control (Cont) or CytoD using sharpened cylindrical probes. Arrows indicate where the cylindrical probe penetrates the cell (“Indent-site”). Scale bar represents 5 µm. **C, D)** Change in lifetime at the ER and nucleus membrane (Nuc-mem) during (C, noted as “mech + inj” in B) and after (D) stimuli imposing injection of DMSO as control (light) or CytoD (dark) at different subcellular compartments: nucleus (top panel), ER (middle panel) and cell periphery (bottom panel) (Mean ± SD, N = 2, n ≥ 5, **p < 0.01, *p < 0.05, t-test). Orange arrows in C indicate the injection pulse at the specified time in the mechanochemical process. Mean values are shown with triangles in boxplots in D (N = 2, n ≥ 5, *p < 0.05, reported p-values of 0.05 ≤ p < 0.1, ns: not significant, t-test).

Higher tension at the ER for control cells compared to CytoD-injected cells suggests actin networks transmit strain and tension intracellularly, unlike intermediate filaments which absorb mechanical stress, preventing it from reaching the nucleus.^40^ Therefore, interrupting actin filaments likely keep tension localized at the indentation site. This is validated by higher tension transfer to the ER in control cells compared to CytoD-treated ones when the cell periphery is targeted (Fig. 3C, top and bottom rows). The ER’s complex morphology consists of many curved structures of lumens and disks which can deform when subjected to external mechanical perturbation, making it prone to absorb stress and strain when directly targeted. That could explain the similar tension levels in both CytoD-injected and control cells during ER indentation. Notably, unlike nucleus-targeted cells, ER-targeted cells only show increased tension at the ER and NE in CytoD-injected cells compared to control ones after stimulation. This suggests fundamental differences in the mechanical coupling of the nucleus and the ER to the cytoskeleton. While the nucleus maintains direct, structural actin connections through the nuclear lamina and LINC complexes^41^, making it immediately sensitive to actin disruption, the ER relies on indirect, signaling-mediated cytoskeletal interactions^42,43^. Thus, the ER functions as a dynamic tension buffer that engages cytoskeletal support primarily during active repair processes following mechanical perturbation. This explains why nuclear probing reveals immediate tension differences in actin-disrupted cells, while ER probing only shows post-retraction effects. Consequently, statistical analysis detects significant differences between test and control groups after probe removal, but not during probe contact in ER-targeted experiments. Yet, when the nucleus is targeted, higher tension at both the NE and the ER is recorded for CytoD-treated compared to control cells both during and after the stimuli. Such distinct mechanoresponse patterns from the ER and the NE (Fig. 3C, top and middle rows) also hint at a directional mechano-crosstalk between the ER and the nucleus of a stimulated cell.

Nuclear lamins regulate nuclear mechanoresponse. Therefore, we investigated how different lamins impact mechano-signaling in HeLa cells upon mechano-chemical stimulation of the nucleus, to shed light on dynamic tension responses.

### Lamin A/C’s autonomous contribution to nucleus mechanoresponsiveness and its dependence on B-type lamins only beyond a sustained force threshold

The nuclear lamina regulates nuclear tension and contributes to cellular mechanotransmission.^44^ Lamin A/C modulates nuclear stiffness and chromatin remodeling,^11^ while its spatial distribution determines nuclear deformability.^45^ B-type lamins have little effect on nuclear stiffness^45^ but maintain NE integrity.^46^ Moreover, a balance between lamin A/C and lamins B1 and B2 is important for nuclear elasticity, with an increased ratio of lamin A/C to the B-type lamins corresponding to a stiffer nucleus. The interplay between A- and B-type lamins, their post-translational modifications, and their impact on governing the nuclear deformability thus genome organization remain underexplored.^47^ Here, we examined the contribution of nuclear lamins to changes in membrane tension using HeLa cells with altered nuclear lamina composition upon downregulation of A-type and/or B-type lamins.

Firstly, WT and *LMNA* KO HeLa cells were subjected to RNA interference (RNAi), using a non-targeting silencing RNA (siRNA) control pool (siCtrl) or pools of siRNAs targeting lamin B1 and lamin B2 (siLMNB) for simultaneous KD of both B-type lamins. The use of RNAi yielded 4 groups of conditions: WT-siCtrl (WT), *LMNA* KO-siCtrl (A-KO), WT-siLMNB (B-KD) and *LMNA* KO-siLMNB (A-KO/B-KD). Western blot analysis confirmed successful depletion of corresponding lamins in each group (*SI Appendix*, Fig. S4). Cells were then stained with the ER Flipper TR dye and assessed for changes in tension levels (Fig. 4A). Lifetime values at the ER and NE were quantified and compared for the different groups (WT, A-KO, B-KD and A-KO/B-KD; Fig. 4B). Higher tension was captured at the ER and NE for both A-KO and B-KD compared to WT cells. A-KO cells displayed significantly higher tension at the ER compared to WT cells, whereas B-KD cells showed a slightly elevated tension level in comparison to WT. Surprisingly, the A-KO/B-KD group showed no captured changes in tension at either the ER or NE in comparison to WT cells. Next, we indented cells by cylindrical probes followed by CytoD injection (Fig. 4C), and quantified tension level in the same manner as before (Fig. 4 D). A significantly lower tension at the ER, NE and indentation site for A-KO compared to WT cells were detected. Intriguingly for B-KD cells, similar tension levels were observed at both the NE and the indentation site when compared to WT cells. At the ER though, like A-KO cells, B-KD cells showed lower tension in response to external stimulus when compared to WT cells. Surprisingly, no changes in tension level of A-KO/B-KD cells were detected compared to WT cells. As varying lamin compositions may alter nuclear stiffness, we also measured indentation depth under a constant 150 nN force (SI Appendix, Fig. S6). We found no statistically significant difference in the indentation depth between groups. The differing tension responses are therefore due to intrinsic differences in molecular mechanics under identical stimulus. Examining the kinetic traits of WT and A-KO/B-KD cells revealed a reduced slope in the tension response from cells lacking A-type and B-type lamins, particularly noticeable in the steady-state response at 4 min post-stimulus (*SI Appendix*, Fig. S7).

**Fig. 4.**
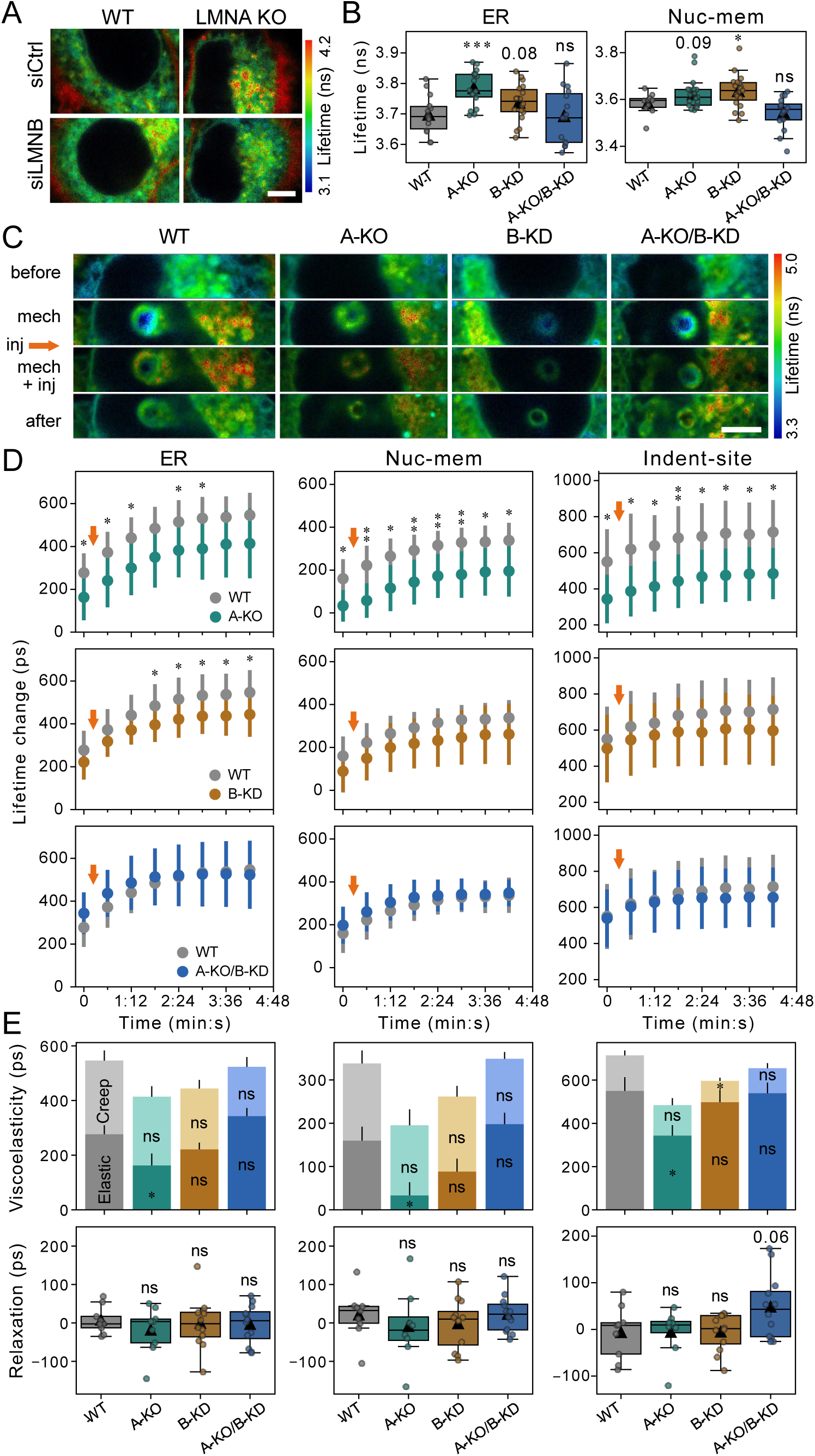
A-type lamins contribute to elasticity, while B-type lamins may influence viscosity in the nucleus’s viscoelastic behavior under sustained deformation with disrupted actin polymerization. **A, B)** Lifetime images (A) and boxplots showing the absolute lifetime values (B) of HeLa cells with varied levels of downregulation of lamin proteins: WT-siCtrl (WT), WT-siLMNB (A-KO), LMNA KO-siLMNB (B-KD) and LMNA KO-siLMNB (A-KO/B-KD). Cells were stained with ER Flipper-TR dye. Scale bar in A represents 5 µm. Mean values are shown with triangles in B (N = 2, n ≥ 8, ***p < 0.001, *p < 0.05, reported p-values of 0.05 ≤ p < 0.1, ns: not significant, t-test). **C)** Lifetime images of WT, A-KO, B-KD and A-KO/B-KD cells stained with ER Flipper-TR before, upon indentation (“mech”), upon indentation + CytoD injection, “mech + inj”, and after stimulation using sharpened cylindrical probes. Note the extended range for lifetime, compared to A, revealing distinguished cell responses to the stimuli, thus indicating the initial tension state for different cell versions in the same color. Scale bar represents 5 µm. **D)** Lifetime change at the ER, nucleus membrane (Nuc-mem) and indentation site during mechanochemical stimuli. All cells were injected with CytoD. In each panel, the tension response of WT cells were compared with either of A-KO, B-KD and A-KO/B-KD cells. Orange arrows indicate the injection pulse at the specified time in the mechanochemical process. Time zero corresponds to the indented state (indicated by “mech” in C) which follows by an injection (indicated by “mech + inj” in C) (Mean ± SD, N = 3, n ≥ 9). **E, top)** Viscoelastic response of cell compartments (left to right: ER, Nuc-mem and Indent-site) with varied depletion of lamins to the mechanochemical stimuli composed of lifetime change originating from the instantaneous cell response (Elastic) upon indentation in nucleus (first time points in D) and from the time- dependent response (Creep) due to AFM probe being in contact with the nucleus (at 150 nN) for around 4 min after CytoD injection (last time points in D). Data represent the mean ± SEM (N = 3, n ≥ 9, ***p < 0.001, **p < 0.01, *p < 0.05, reported p-values of 0.05 ≤ p < 0.1, t-test). **E, bottom)** Relaxation behavior of WT, A- KO, B-KD and A-KO/B-KD cells at varied positions (left to right: ER, Nuc-mem and Indent-site) after stimuli reported as lifetime change right after probe retraction compared to last time point during stimuli (last time points in D). Mean values are shown with triangles in boxplots (N = 3, n ≥ 9, reported p-values of 0.05 ≤ p < 0.1, ns: not significant, t-test).

These results align with earlier findings stating that cells lacking lamin A/C, but not Lamin B1 and B2, have softer nuclei with a more deformable NE, thus dissipating the stress and lowering the tension at the NE^48,49^ This leads to a lower tension level at the NE for A-KO cells compared to WT cells when exposed to external stimuli. The elevated wrinkles in the NE of lamin A/C KO cells^50^ might play a role in dissipating stress and reducing tension during mechanical stimuli, particularly when these wrinkles open due to the deformation. In essence, a nucleus without A-type lamins appears less mechanoresponsive, as it likely absorbs forces through bending rather than transferring them. In agreement with the previous studies, B-type lamins on the other hand, do not affect nuclear stiffness, and deformation, thus its tension response.

Lower tension in A-KO and B-KD cells compared to WT cells suggests that both A-type and B-type lamins influence the tension crosstalk from the NE to the ER. Examining their tension response reveals that when actin filaments are disturbed, lamin A/C primarily affects the immediate response of nucleus to physical deformation of the nucleus. Conversely, B-type lamins become more significant under sustained load. In other words, lamin A/C alone can contribute to the nucleus’s mechanoresponsive abilities, but *LMNA* KO cells become less mechanoresponsive than WT cells and depend on the supportive function of B-type lamins. This implies that even at this juncture, A-KO/B-KD cells under prolonged stress might show reduced mechanoresponses compared to WT cells, notably with a decrease in tension at the indentation site recorded after 3 minutes post-stimulus (Fig. 4D).

The nucleus stretches rapidly in response to a step-function load and undergoes a viscoelastic transition with sustained load, taking seconds to reach a steady state. This behavior depends on lamina composition and chromatin states,^51^ possibly aiming to dissipate mechanical energy and prevent nuclear damage. We discerned the different roles of lamin A/C versus lamin B1 and B2 in the viscoelastic response of the nucleus and the ER. This process separated the immediate response (“elastic”), due to the probe indentation measured at “mech” state from a gradual deformation (“creep”) under constant force applied during “mech + inj” state. Creep response reflects cell’s viscous property, with higher creep reflecting lower cell’s viscosity. The viscoelastic response was influenced by CytoD injection as a chemical stimulus, followed by a continuous load of 150 nN for 4 min ± 10 s. The results are depicted as accumulated bar plots in the top panel of Fig. 4E. At the ER, similar levels of elastic and viscous responses were detected for cells with different lamin proteins levels compared to WT cells. At the NE, though, lower elasticity in *LMNA* KO cells and similar viscous response in all the cells were captured. At indentation site, cells lacking A-type lamins show a reduced elastic property while cells with deficiencies in B-type-lamins have lower creep and thus higher viscous response. Further the temporal relaxation of tension at the ER, NE and indentation site were investigated both immediately after the stimulus (Fig. 4E, bottom row) and 5 min post-stimulus (*SI Appendix*, Fig. S8). Relaxation property was quantified by comparing the lifetime value after probe retraction to the last time point of stimuli at “mech + inj” state. A higher relaxation value indicates greater recovery of the deformed structure and thus a stronger elastic component of the mechanoresponse. Following the stimulus, cells deficient in both A-type and B-type lamins displayed increased relaxation at the indentation site which experiences maximum deformation (Fig. 4E, bottom right). Finally, 5 min post-stimulus, cells with diverse depletion of lamins exhibit a consistent relaxation extent across various ROIs (*SI Appendix*, Fig. S8). These findings suggest A-type lamins contribute to nuclear elasticity, while B-type lamins influence viscous behavior during sustained deformation and disrupted actin polymerization (Fig. 4E, top right). Nonetheless, previously a potential role for lamin A in viscous response of NE in both intact tissue and suspended cells has been reported.^52,53^ In our study, the disruption of the dynamic actin network by CytoD injection suggests that the interaction between lamin A/C and the actin meshwork may be essential for maintaining the viscous properties of lamin A. Higher relaxation in A-KO/B-KO cells immediately upon removal of probes, reflecting higher tension restoration, may be linked to the absence of actin filaments alongside the lamins depletion.

In contrast, WT cells and cells lacking either A-type- or B-type-lamins maintain tension, releasing it gradually over time. Similar relaxation values for all cells at 5 min post-stimulus indicate cytoskeletal reorganization and NE adaptations, allowing stress recovery independently of nuclear lamina. Since nuclear lamina organization affects how cells detect and respond to mechanical stimuli, we aimed to dissect the roles of cytoskeletal components in initiating the mechanoresponse of the nucleus and ER after mechanical and chemical stimuli with various molecular disruptors (CytoD, Noco, and their combination).

### Modulating mechanoresponse: impact of lamin A/C on microtubule dynamics and adaptive mechanosensitivity

The nuclear lamina contributes to organize the cytoskeleton^40^, with other components like actin filaments and microtubules enabling cells to sense and respond to mechanical stimuli (Fig. 2G).^54^ In particular, loss of lamin A/C is linked to decreased myosin II contractility and in some cases to the absence of apical stress fibers,^55^ which disrupts cytoskeletal dynamics and affects how cells interpret mechanical signals. Conversely, MT depolymerization can enhance actomyosin contractility via the RhoA-Rock pathway, altering cell mechanics and nuclear properties. Considering the intricate interplay between the cytoskeleton, nucleus, and mechanical cues in cellular behavior, we assessed how changes in cytoskeletal components and lamin A/C affect cellular mechanoresponse and disease contribution.

Noco- and CytoD-Noco-treated WT cells had low tension across ROIs. A-KO cells showed higher tension at the ER and NE with Noco or CytoD-Noco compared to CytoD alone. No difference in tension between Noco and CytoD-Noco groups at the ER and NE.

Actomyosin and microtubules, due to their dynamic response compared to intermediate filaments^56^, were studied for their roles in initiating nuclear mechanoresponse. We targeted the cytoskeleton of WT and A-KO cells with different disruptors including CytoD, Nocodazole (Noco, a microtubule inhibitor), and their combination (CytoD-Noco), each at 50 µM, resulting in 0.5 µM upon intracellular injection. Noco binds to β-tubulin subunits, thereby inhibiting the polymerization of tubulins into MTs.^57^ These investigations focused on WT and lamin A/C deficient cells, due to lamin A/C’s distinctive roles in NE mechanoresponse as examined in section 2.4. Tension response to stimuli were quantified at various ROIs (Fig. 5 A and B). Notably, WT cells mechanoresponse is governed by MTs presence, showing increased tension in CytoD-treated cells compared to Noco- or CytoD-Noco-treated cells at the ER, NE, and the indentation site. Tension in WT cells with MTs (CytoD-treated cells) compared to those with less MTs (either Noco- or CytoD-Noco-treated cells) significantly increased upon stimulation. Noco- and CytoD-Noco-treated WT cells exhibited similarly low levels of tension across all ROIs. Cells lacking lamin A/C showed distinct responses to the aforementioned drugs compared to WT cells. *LMNA* KO cells displayed higher tension at the ER and NE when subjected to Noco or CytoD-Noco (less MTs in function) compared to when exposed to CytoD alone. Interestingly, at the ER and NE there were no difference in tension values between Noco and CytoD-Noco groups. At the indentation site, A-KO cells showed similar trends in the ER and NE for CytoD and Noco treated cells. Interestingly, A-KO cells lacking both actin filaments and MTs had the lowest tension.

**Fig. 5.**
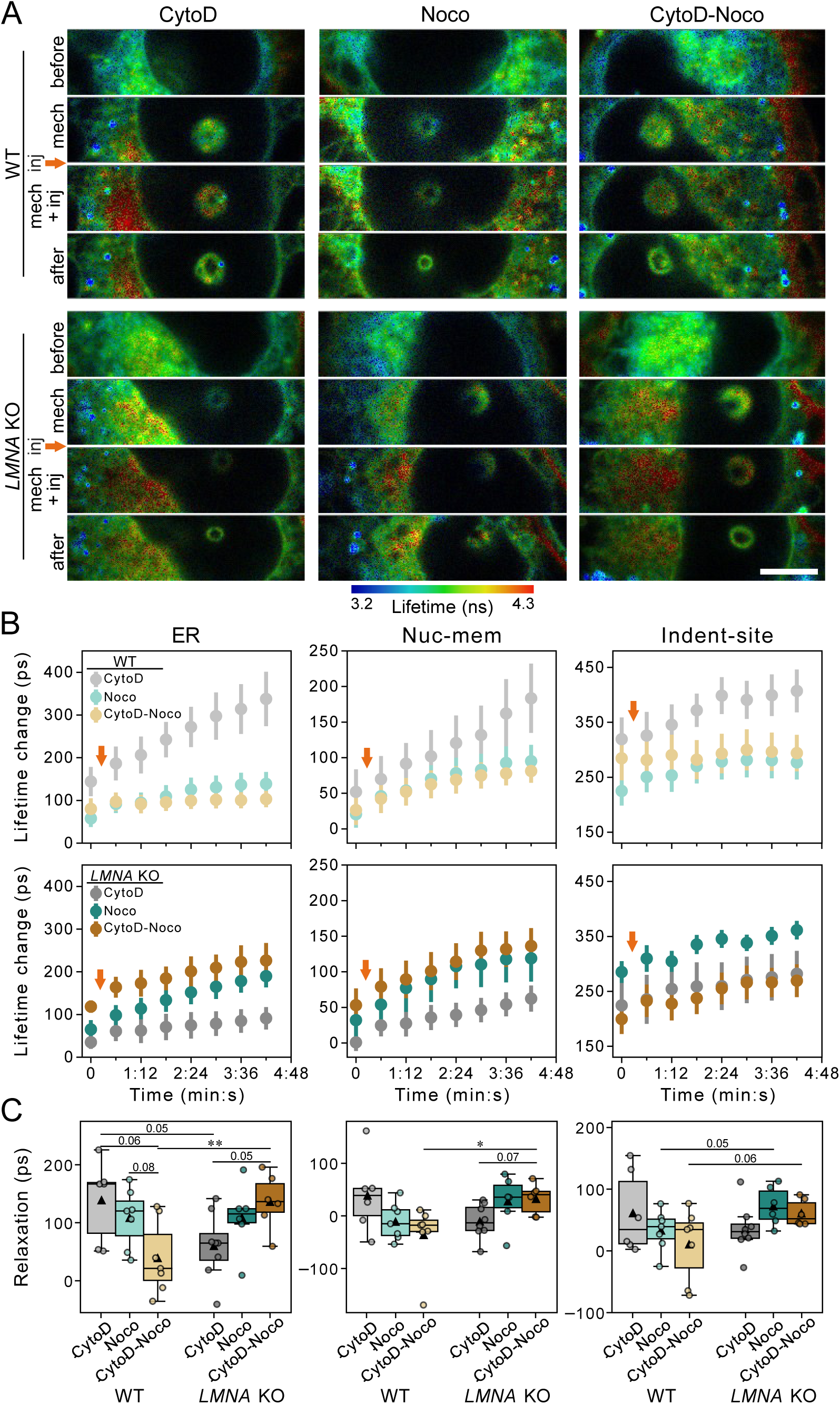
Dynamic role of microtubules and actinomyosin in regulating mechanoresponse of nuclei in cells with and without lamin A/C-depletion to the external stimulus. **A)** Lifetime images of WT- and LMNA KO-HeLa cells stained with ER Flipper-TR before, upon indentation (“mech”), upon indentation + injection (“mech + inj”) of various drugs and after stimulation using sharpened cylindrical probes. For chemical stimulation, interruption of different cytoskeleton components including actin filaments, microtubules and both of actin filaments and microtubules was targeted realized by injection of 50 µM CytoD (left column), 50 µM Noco (middle column) and 50µM CytoD + 50µM Noco (right column) dissolved in DMSO. Scale bar represents 5 µm. **B)** Lifetime change at ER, nucleus membrane (Nuc-mem) and indentation site during stimuli in WT and LMNA KO cells. Arrows indicate the injection pulse at the specified time in the mechanochemical process. Time zero corresponds to indented state (indicated by “mech” in A) which follows by injection, indicated by “mech + inj” in A (Mean ± SEM, N = 2, n ≥ 6, **p <0.01, *p < 0.05, reported p-values of 0.05 ≤ p < 0.1, t-test with Holm correction). **C)** Relaxation behavior of WT and LMNA KO cells at varied positions (left to right: ER, Nuc-mem and Indent- site) 5 min after mechanochemical stimuli involving various drugs (CytoD, Noco and CytoD-Noco). Relaxation is reported as lifetime change at 5 min after probe retraction compared to the last time point of stimuli (last time points in B). Mean values are shown with triangles in boxplots (N = 2, n ≥ 6, **p <0.01, *p < 0.05, reported p-values of 0.05 ≤ p < 0.1, t-test with Holm correction).

Higher tension in WT cells with active MTs compared to those with less MTs (Noco- and CytoD-Noco-treated) indicates MTs’ dominant role in tension propagation within the ER-NE network. This is likely due to their close association with the ER network. Microtubules seemingly maintain tension within the ER-NE network rather than transferring it, independently of actin filaments (Fig. 5B, top row). Yet, actin filaments lower the tension in cells with intact MTs (Fig. 2 E and F). In A-KO cells, MTs facilitate nuclear deformation.^58^ Thus, in A-KO cells, Noco and CytoD-Noco treatment minimizes the deformation of nucleus and increases tension at the ER and NE. On the other hand, CytoD treated A-KO cells with functional MTs exhibit higher deformation and thus lower tension. Similar tension levels in Noco- and CytoD-Noco-treated A-KO cells at the ER and NE indicate actin filaments independence. Lamin A/C forms the perinuclear actin cap^50,59^, explaining high tension at the indentation site in Noco-treated A-KO cells compared to WT cells despite having active actin filaments. Low tension in CytoD-Noco-injected A-KO cells suggests a highly deformable nuclear membrane, despite lacking active microtubules at the indentation site.

Next, elastic and viscous (derived from creep behavior) components of tension response in WT and A-KO cells under various drug treatments were quantified (*SI Appendix*, Fig. S9A). WT cells showed decreased creep (increased viscous response) of WT cells with CytoD-Noco compared to CytoD treatment, but no significant difference between CytoD-Noco and Noco treatment across all ROIs. In contrast, A-KO cells had the lowest creep (highest viscous response) in the ER with CytoD treatment. In other words, under lamin A/C deficiency, microtubules reduce tension and diminish the creep response.

Finally, temporal tension relaxation was studied upon probe removal and diminishing the mechanical stimulus. Relaxation was quantified as a change in lifetime immediately after probe retraction (*SI Appendix*, Fig. S9) and 5 min (Fig. 5C) post-stimulus, both with respect to the last time point during stimuli. WT and lamin A/C deficient cells exhibited similar tension immediately after probe retraction, regardless of the injected drug. Additionally, lamin A/C deficient cells treated with various drugs had consistent relaxation compared to WT across all ROIs. Though, relaxation at 5 min post-stimulus depended on drug treatment, especially at the ER and NE. WT cells treatment with CytoD or Noco alone had higher relaxation compared to CytoD-Noco at the ER. Instead, A-KO cells showed the lowest relaxation with CytoD and highest with CytoD-Noco treatment, 5 min post stimulus at the ER and NE. Notably, no significant differences in relaxation were found among drug treatments at the indentation site for both WT and *LMNA* KO cells.

Collectively, both actin and microtubules contribute to the cell’s elastic mechanoresponse, as shown by the similar elastic relaxation in WT cells treated with CytoD or Noco. Interestingly, loss of both A-type and B-type lamins does not affect tension at the NE or ER, unlike the individual loss of either lamin type. This highlights the complex interplay between different lamins and cytoskeletal elements in cellular mechanics. Microtubules play a lesser role in conferring viscous properties (time-dependent) to healthy cells, indicated by lower creep and higher viscous response of the ER when exposed to microtubule-disturbing drugs (Noco or CytoD-Noco). This is due to the rapid dynamics of microtubules with critical roles in cell division and intracellular transport.^60,61^ In lamin A/C-deficient cells, MTs contribute to a more viscous mechanoresponse, shown by higher creep and lower viscous response with microtubule-disturbing drugs (Noco or CytoD-Noco). This adaptation in microtubule mechanosensitivity may be due to the absence of the actin cap in lamin A/C-depleted cells.

## Conclusion

Combining FluidFM with FLIM enabled real-time, high-resolution mapping of intracellular tension dynamics upon quantitative injection of active components while maintaining a constant mechanical stimulus. The proposed approach allowed investigation of intracellular mechanotransmission crucial for mechanotransduction regulation and cell fate control. CytoD Injection, as *in situ* mechano-chemical trigger, disrupted actin filament dynamics, altering tension at the ER and NE. Sharpened cylindrical probes increased tension locally on the nucleus, and CytoD-injection increased tension at the ER and NE compared to control injection. Pyramidal probes required a higher force to puncture the nuclear membrane (50 nM) compared to that of the cylindrical one (10 nN). Membrane puncture resulted in tension relaxation at the NE. Increased indentation force for sealed injection showed lower tension at the NE and the ER with cylindrical probes compared to pyramidal ones. Changing the incident site from the nucleus to the ER and cell periphery confirmed that actin filaments alone transmit tension away from the deformation site. In contrast, CytoD injection and actin filament disturbance retained tension at the site of incidence. Direct ER targeting suppressed tension from reaching the nucleus. This indicates specialized tension distribution within cells. Nuclear lamins showed distinct responses to mechanical cues, with A-type and B-type lamins affecting elastic and viscous properties, respectively. Interestingly, combined loss of lamins A and B did not affect tension at the NE or ER, suggesting a compensatory mechanism needing further investigation.

In conclusion, our study examined how the cytoskeleton, particularly actin and microtubules, collaborates with nuclear lamina to initiate nuclear mechanoresponse. In WT cells MTs maintained tension, while actin filaments lowered it in responding to external stress. In lamin A/C-deficient cells, microtubules facilitated nuclear deformation and lowered the tension, in contrast to actin filaments. This highlights lamin A/C’s role in regulating the collaboration between the NE and the cytoskeleton, affecting nuclear and the ER mechanoresponse. Additionally, nuclear deformation under external stimulus involves interactions between chromatin and cytoskeletal components through the lamina and LINC complexes. However, the exact mechanisms of chromatin involvement require further study. Lastly, the intermediate filament system, particularly vimentin, also plays a key role in nuclear integrity and connecting to other organelles like mitochondria. Future studies should explore their contributions to mechanotransmission phenomena.

## Methods

For detailed procedures of the following steps, see *SI Appendix*.

### Cells Preparation

HeLa cells with WT condition or with altered levels of lamin proteins were cultured using standard procedure for adherent cells. Cells were seeded on glass bottom dishes 1 or 2 days before the experiments. Just before the experiments, cells medium was changed to CO_2_ independent medium and cells were stained with 1 µM ER Flipper-TR, designed to report changes in membrane tension at the ER. However, ER Flipper-TR dye apparently stains other endomembrane structures in addition to ER as well.

### FluidFM Probe Preparation

The custom-fabricated cylindrical probes (SmartTip BV in Enschede, the Netherlands) featured a tube length of 10 µm, an aperture diameter of 1 µm, and a microchannel thickness of 1.5 µm.^62^ Subsequently, they underwent milling with an FIB machine mounted on a SEM (Helios 5 UX, ScopeM, ETH Zurich, Switzerland) to achieve sharpness at their apex (50° angle, Fig. 1 B). Both customized cylindrical and purchased pyramidal probes (Cytosurge AG, Switzerland) were subjected to coating with Sigmacote vapor to minimize contamination during experiments. This coating imparted anti-fouling properties. The probes were baked at 100°C for 2 hours to ensure stability. Finally, the probes, filled with 0.1 mg·mL⁻¹ coumarin 120 (blue dye) plus the drug of interest (50 µM CytoD, 50 µM Noco, or 50 µM CytoD + 50 µM Noco) in HEPES2 buffer, were used for intracellular injection experiments upon calibration.

### FluidFM-FLIM Setup

We combined FluidFM system (FlexAFM-near-infrared scan head with a C3000 controller driven by the EasyScan2 software, Nanosurf AG, Switzerland) with FLIM imaging on a Leica SP8 FALCON inverted confocal microscope.^31^ To achieve this, we removed the condenser lens and z-galvo stage. The FluidFM probe was independently positioned using an elevated AFM scan head attached to a custom mount. Adjustments were made to reach the substrate surface, and FLIM imaging utilized laser pulses at 488 nm operating at a frequency of 40 MHz. The entire setup was maintained at a temperature of 37°C. The AFM piezo movement, deflection of the cantilever, and fluidic pressure (in the case of injection in mechano-chemical stimulation) were recorded utilizing a data acquisition box and a LabView script.

### Mechanical and Mechano-chemical Stimulation

For the FluidFM stimuli, the probes were first calibrated for spring constant and sensitivity, then approached on the HeLa cells stained with the ER-Flipper-TR dye. The setpoint for the indentation force varied between 2 and 100 nN for mechanical stimulation, while it was adjusted to 150 nN for the mechano-chemical stimulation assuring a successful injection. Both stimuli were operated at an approach speed of 1 µm.s^-1^, which was chosen to mimic the natural mechanical environment that cells experience in the body. For mechano-chemical stimulus employing an injection pulse (100 mbar for 10 s) after reaching the setpoint, probes were previously filled with the desired chemical solution. Lifetime images of the target cells were concurrently captured, featuring a pixel size of 90 nm. For comparison, lifetime images were collected before the stimulus, during the pause time of the mechanical stimulus before injection pulse (“mech”), series of images after the injection pulse (“mech + inj”) and one image after retracting the probe. For experiments addressing the relaxation behavior, an additional lifetime image was captured 5 minutes post-stimulus.

### Statistical Analysis

Data were tested for normal distribution using the Shapiro-Wilk test before applying parametric statistical methods. Differences between two groups were assessed using an unpaired two-tailed Student’s t-test. Where multiple comparisons were conducted, a t-test with Holm correction was applied to control the false discovery rates, as indicated in the relevant sections. P-values were calculated and reported for all analyses, with a significance threshold set at p < 0.05. The sample size for each experiment (n) is provided in the figure legends, and the number of independent biological replicates (N) is explicitly reported to ensure reproducibility. All statistical analyses were performed custom Python scripts.

## Supporting information

all supplementary information

## Acknowledgments

The authors thank János Vörös and all the LBB team for insightful discussions, Edin Sarajlic (Smartip BV, NL) for fabrication of customized FluidFM probes, Stephen Wheeler for his workshop support, Joakim Reuteler and Justine Kusch at ScopeM facility (ETH Zurich) for microscopy support, and Aldo Riccardo at cleanroom facility (D-ITET ETH Zurich). E.Z.E. and I.L. received support from the EUREKA Network (Eurostars Project E!11644 “SOUL” to T.Z.). Additionally, E.Z.E. acknowledge funding from the Swiss National Science Foundation (Sinergia Grant CRSII5_202301 to T.Z.). U.K. acknowledges support by the Swiss National Science Foundation (project grant 310030_219203 and SNSF Advanced Grant TMAG-3_209245).

## Data Availability

The datasets generated and analyzed during the current study, including the FLIM image data and the Python scripts developed for the quantification of values, statistical analysis, and data visualization are available in the ETH Zürich Research Collection at https://doi.org/10.3929/ethz-c-000790250. Additionally, the force and pressure measurements, are available upon request.

## Notes

https://doi.org/10.3929/ethz-c-000790250

### Competing Interest Statement

The authors have declared no competing interest.

### Summary of Updates

Section "Results and Discussion": - details about insertion of pyramidal probes; - hypothesis about tension levels in both CytoD-injected and control cells during ER indentation; - details about laminin indentation depth at constant force setpoint.

