## Supplementary material for "Quantifying Intracellular Mechanosensitive Response upon Spatially Defined Mechano-Chemical Triggering": all supplementary information

**Materials.** DMEM (Dulbecco's Modified Eagles Medium + GlutaMAX, Thermo Fisher Scientific, Waltham, USA), $CO_2$ Independent Medium (Thermo Fisher Scientific, Waltham, USA), FBS (fetal bovine serum, Thermo Fisher Scientific, Waltham, USA), penicillin-streptomycin (Thermo Fisher Scientific, Waltham, USA), Trypsin (Thermo Fisher Scientific, Waltham, USA), PBS (phosphate buffered saline, Thermo Fisher Scientific, Waltham, USA), coumarin 120 (7-amino-4-methylcoumarin, Sigma-Aldrich, St. Louis, USA), SL2 Sigmacote (Sigma-Aldrich, St. Louis, USA), ER Flipper-TR (LubioScience, Zurich, Switzerland), NucBlue (Thermo Fisher Scientific, Waltham, USA)

**Cell Preparation.** HeLa cells were chosen as a model system for mechanoresponse studies due to the availability of genetically modified versions with altered levels of lamin proteins. HeLa cells were cultured at 37°C and 5% $CO_2$ in DMEM culture medium supplemented with 10% FBS and 1% Penicillin-Streptomycin using standard procedure for adherent cells. For splitting, 0.05% trypsin was used. Cells were seeded on glass bottom dishes (Willco, Amsterdam, Netherlands) with a seeding density of 6'000 cells/ $cm^2$ or 4'000 cells/ $cm^2$, 1 or 2 days before the experiments. Just before the experiments, medium was changed to $CO_2$ Independent Medium and cells were stained with 1 µM ER Flipper-TR.

**Generation of lamin knockout (KO) and knockdown (KD) cells**. HeLa lamin a gene (*LMNA*) KO cells have been described previously.[1] RNA interference (RNAi)-mediated depletion of B-type lamins was accomplished by siPOOLs (siTOOLs Biotech). Wild-type HeLa (WT) or HeLa *LMNA* KO cell seeded into 6-well plates and transfected with 5 nM small interfering RNA (siRNA) pool targeting each protein of interest, or a pool of non-targeting control siRNAs using the transfection reagent RNAiMAX (ThermoFischer Scientific, Waltham, USA). After 72 h, cells were harvested for western blots or subjected microscopic analysis. Additionally, to rule out clonal anomalies in the *LMNA* KO cells, HeLa WT cells were knocked down for lamin A/C, using a 5 nM pool of siRNA Subsequently, these *LMNA* KD cells, *LMNA* KO cells and WT cells were stained with ER Flipper-TR and their initial lifetime time values were quantified and compared (Fig. S5).

**Immunoblotting.** Harvested cell pellets were lysed in SDS sample buffer (75 mM Tris pH 7.8, 20% (v/v) glycerol, 4% (w/v) SDS, 50 mM DTT, 0.01% (w/v) bromophenol blue) to yield whole cell lysates. Proteins were separated on pre-cast SDS-PAGE gels (vendor), and then transferred to a nitrocellulose membrane (GE Healthcare) using semi-dry blotting. The membrane was then blocked using blocking solution (5% milk in PBST) for 30 min, before being incubated with primary antibody diluted in blocking solution overnight at 4°C. The membrane was washed thrice for 5 min using blocking solution and subsequently incubated with secondary antibody diluted in blocking solution. Finally, the membrane was washed three times using PBST. Membranes were then developed using a LI-COR (Biosciences) to probe for a protein of interest.

**FluidFM Probe Preparation.** As detailed previously[2], cylindrical probes were custom-fabricated (SmartTip BV, Enschede, the Netherlands) featuring a tube length of 10 µm, an aperture diameter of 1 µm, and a microchannel thickness of 1.5 µm. Cylindrical probes were affixed to a custom-made probe holder for carbon coating (18 nm) using a CCU-010 Carbon Coater (Safematic, Switzerland) before milling with an FIB machine mounted on a SEM (Helios 5 UX, ScopeM, ETH Zurich, Switzerland) to achieve sharpness at their apex at a 50° angle (Fig. 1 B). FIB-milled cylindrical probes were then glued onto a cytoclip holder by Cytosurge (Cytosurge AG, Switzerland). Pyramidal probes (Nanosyringes, Cytosurge AG, Opfikon, Switzerland) and cylindrical probes were both subjected to oxygen plasma treatment (100-E Plasma System, Technics Plasma GmbH, Munich, Germany) for 2 minutes. Following this, they underwent an overnight coating with Sigmacote vapor inside a vacuum-sealed glass desiccator. This coating imparts anti-fouling properties to the probes, minimizing contamination by cell debris during experiments. Subsequently, the probes were baked at 100°C for 2 hours to ensure a stable coating. They were then stored alongside humidity-absorbing salt (Drierite, calcium sulfate, Sigma-Aldrich, St. Louis, USA) and utilized within 3 days post-coating. Prior to conducting the experiments, the spring constant of empty probes, initially designated with a nominal value of 1 $N.m^{-1}$, was determined using optical beam deflection and employing the Sader method.[3,4] Subsequently, the deflection sensitivity of the probes, filled with 0.1 $mg.mL^{-1}$ coumarin 120 (blue dye) plus the drug of interest for intracellular injection (50 µM cytochalasin D (CytoD), 50 µM nocodazole (Noco), or 50 µM CytoD + 50 µM Noco) in HEPES2 buffer, was measured.

**FluidFM-FLIM Setup.** As detailed previously[5], FluidFM setup for micromanipulation of cells combined with FLIM imaging setup on the Leica SP8 FALCON inverted confocal microscope (Leica Microsystems GmbH, Wetzlar, Germany). To facilitate This, the condenser lens and z-galvo stage of confocal microscope were removed. For independent positioning of the FluidFM probe with respect to cell dish, the AFM scan head (Nanosurf AG, Liestal, Switzerland) was elevated above the standard microscopy stage using a custom mount attached to the microscope's incubator box. This mount, with an integrated manual x-y-positioning system, hovered approximately 0.5 mm above the microscopy stage. To reach the substrate surface despite the elevated AFM scanhead, adjustments were made to the z-screw holders and z-motor housing, and magnets were placed to lower the FluidFM probe by around 5 mm. The cell dish was positioned on a spaced sample-holder, allowing access for microscopy using a 63x oil objective with a numerical aperture of 1.4 (HC PL APO CS2, Leica Microsystems GmbH, Wetzlar, Germany). AFM laser alignment was monitored using an ocular camera (DinoEye, AnMo Electronics Co, Taipei, Taiwan). FLIM imaging utilized laser (488 nm) pulses operating at a frequency of 40 MHz. The microscope chamber was maintained at a temperature of 37°C. The AFM piezo movement, deflection of the cantilever, and fluidic pressure (in the case of injection in mechanochemical stimulation) were tracked by recording timelines using a data acquisition box and a custom LabView script (National Instruments Co, Austin, USA). These timelines were then correlated with the recorded lifetime images through a trigger output from the confocal microscope.

**Mechanical stimulation.** For the mechanical stimulation, FluidFM probes were approached on the ER Flipper-TR-stained HeLa cells. The set point for the indentation force varied between 2 and 100 nN, while the approach speed was set at 1 $\mu m.s^{-1}$. Simultaneously, lifetime images of the target cell were captured with a field of view measuring 92.35 μm x 23.09 μm, featuring a pixel size of 90 nm. These images were scanned at a speed of 100 Hz, utilizing a pinhole aperture of 1.0 airy unit (equivalent to 95.5 μm). Photons from 30 frames per image were collected and averaged, with an acquisition time of 40 s for each image. For comparison, one image was taken before the mechanical stimuli, one during the pause time of the stimulus, and one after removing the probe.

**Mechanochemical stimulation.** For mechanochemical stimulation, FluidFM probes filled with the desired chemical solution were approached on the ER Flipper-TR-stained HeLa cells with varied levels of lamin proteins. The setpoint was adjusted to 150 nN for the indentation to ensure a complete insertion of the probe aperture inside the cell, thereby a sealed injection, with an approach speed of 1 $\mu m.s^{-1}$. After reaching the setpoint and undergoing a brief waiting period, an injection pulse of 100 mbar for 10 s was administered. The injection parameters, comprising the indentation force, approach speed, and fluidic pressure pulse, had been specifically calibrated for optimal application to HeLa cells beforehand. Concurrently, lifetime images of the target cell were acquired for up to 5 min post-injection, covering a field of view spanning 46.13 μm x 5.69 μm and featuring a pixel size of 90 nm. These images underwent scanning at a rate of 200 Hz, utilizing a pinhole aperture of 1.0 airy unit (equivalent to 95.5 μm). Each image was composed of photons collected from 100 frames, which were then averaged, with an acquisition time of 18 s for each image. For comparison, lifetime images were captured before the stimulus, during the pause time of the mechanical stimulus before injection pulse ("mech"), series of images after injection pulse ("mech + inj") and one after removing the probe. The chosen observation period of up to 5 minutes post-injection was defined by the expected rapid pharmacokinetics of the drugs upon direct intracellular delivery. To achieve this, a high-concentration stock (50 μM) of CytoD and Nocodazole was used to yield a final intracellular concentration of 0.5 μM. This physiologically relevant dose was anticipated to act as fast as within 3 minutes for CytoD and within 5 minutes for Nocodazole, a timing crucial for our study's focus on the early dynamic response and initial deviation from control conditions rather than the steady state achieved at longer time points. This time estimation is firmly based on reported values in the literature. For example, a recent comprehensive study quantified actin dynamics upon CytoD application using TIRF microscopy and reported the onset of actin polymerization inhibition to occur approximately 150 seconds after introducing

5 nM CytoD.[6] Similarly, another study demonstrated the rapid disassembly of microtubules in monocytes incubated with 1 μM Nocodazole, reporting a half-time of 40 seconds for complete disassembly.[7]

For experiments studying the relaxation behavior, an additional lifetime image was captured 5 minutes post-stimulus.

**FLIM Data Analysis.** Firstly, different "Region of Interests" (ROIs) of the endoplasmic reticulum (ER), nuclear membrane (Nuc-mem), and the indentation site (Indent-site) (a region around the indenting probe) were manually selected (Fig. S1). The circular structure enveloping the nucleus as detected in lifetime images of cells stained with ER Flipper-TR (Fig. 1C), was attributed to the ROI associated with the nuclear membrane. To validate this correlation with the nuclear membrane, cells were co-stained with NucBlue (DNA-binding dye) and ER Flipper-TR. The resulting images captured the overlapping signals (Fig. S2). The substantial overlap observed affirms the accuracy of the defined ROI as the nuclear membrane in this study. A two-exponential reconvolution fit was conducted on the time-correlated single-photon counting histogram obtained for every pixel within selected ROIs in all lifetime images. This process allowed for the determination of the two components of the ER Flipper-TR lifetime, with the smaller component being fixed for each image and disregarded for all subsequent analyses. The larger component, which is directly related to membrane tension, was determined for each pixel, resulting in the spatial distribution of membrane tension during mechanical or mechanochemical stimulation. Previously, we verified that changes in larger component of Flipper-TR lifetime during FluidFM stimulation were mainly associated with alterations in tension rather than lipid order.[5] For that, we employed the Laurdan-staining assay, which utilizes Laurdan, a fluorescent dye sensitive to variations in lipid organization within the membrane. By applying a threshold to the fluorescence intensity of each pixel, membranal structures were distinguished from the background. Subsequently, the amplitude-based average lifetime $\tau_{amp}$ was calculated (Eq. 1) for a selected region of interest encompassing the ER, nuclear membrane, and indentation site,

$$\tau_{amp} = \frac{\sum_i A_i \times \tau_i}{\sum_i A_i} \qquad \text{Eq. 1}$$

where $A_i$ represents the amplitude and $\tau_i$ denotes the lifetime for the $i^{th}$ pixel.

The mean lifetime values were determined for the periods before, during, and after the stimulus across all ROIs. Subsequently, the value associated with the pre-stimulus state of each cell was subtracted from its respective image for the during and after periods, resulting in the calculation of lifetime change values. The obtained fluorescence lifetimes are associated with membrane tension rather than lipid composition, as examined in our previous study.[5] However, the conversion factor for ER Flipper-TR were neither addressed previously nor studied here due to the challenges in establishing an experimental setup to measure the ER tension in intact cells. Therefore, the lifetime values are reported directly.

**SI APPENDIX FIGURES**

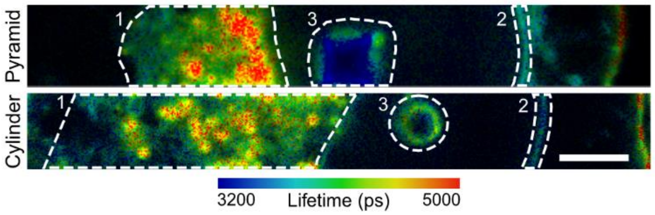


**Fig. S1.** Selection of ROIs of 1) the ER, 2) nuclear membrane (Nuc-mem), and 3) a spot around the indentation site (Indent-site) using pyramidal (top) and cylindrical (bottom) probes. Representative lifetime images of a HeLa cell stained with ER Flipper-TR dye are shown. Scale bar represents 5 µm.

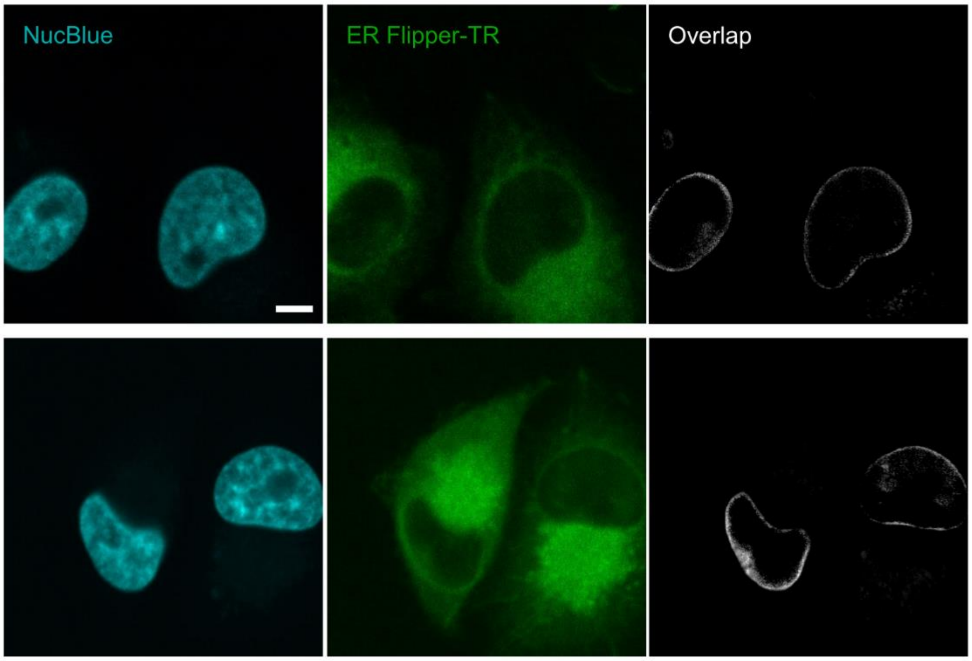


**Fig. S2.** Co-staining HeLa cells with NucBlue (targeting DNA inside the nucleus) and ER Flipper-TR dye. The significant overlap signals obtained provide confirmation of the precision of the designated ROI as representing the nuclear membrane in this investigation. Scale bars represent 5 µm.

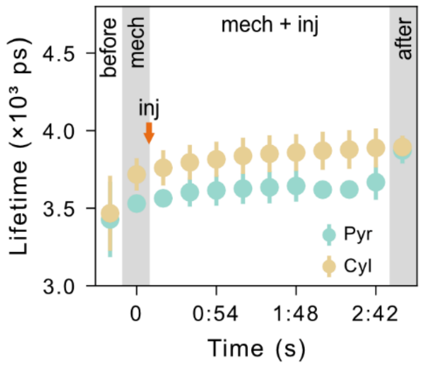


**Fig. S3.** Lifetime values before, during and after injecting HeLa cells with DMSO as control group with pyramidal and cylindrical probes at incident point (Mean ± SD, N ≥ 3, n ≥ 11). Different sections on the graph represent lifetime measurements before stimulus, during mechanical stimulus when the probe has reached the set point on the cell without fluidic pressure application ("mech"), and during mechano-chemical stimulus when the probe remains in contact with the cell at the set point, and a fluidic pressure pulse is applied for drug (or here control) injection ("mech + inj"), and after retraction of probe and removal of the stimulus.

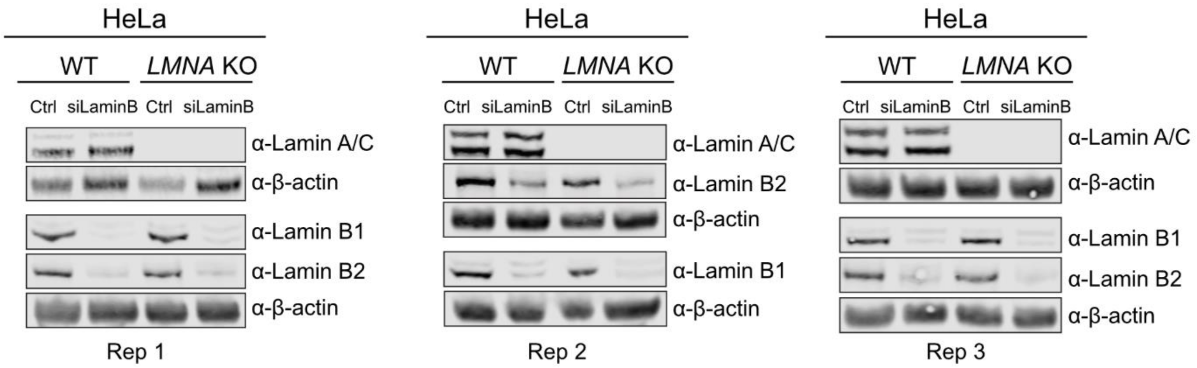


**Fig. S4.** Western blot analysis verifying the absence of lamin A/C in *LMNA* KO cells and the successful downregulation of lamins B1 and B2 in both WT and *LMNA* KO upon treatment with siLMNB. Three biological replicates were utilized for mechano-chemical triggering assays and FLIM-based tension measurements.

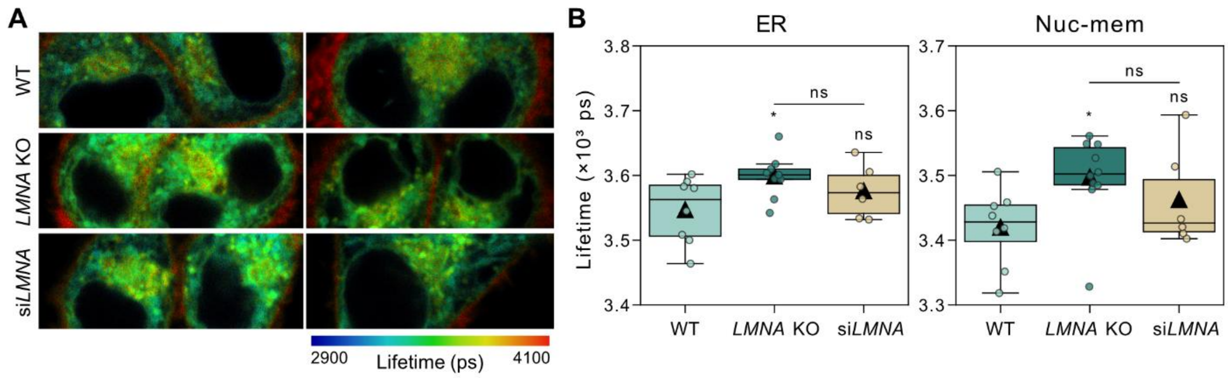


**Fig. S5.** Lifetime images (A) and boxplots showing absolute lifetime values at the ER and nuclear membrane (Nuc-mem) (B) of HeLa cells stained with ER Flipper-TR tension reporter molecule. The lamin A/C level of cells was downregulated with altering methods: WT, *LMNA* KO and WT siLMNA. Cells. Scale bar in A represents 5 µm. Mean values are shown with triangles in boxplots in B (N = 2, n ≥ 6, $^*p < 0.05$, ns: not significant, *t*-test with Holm correction).

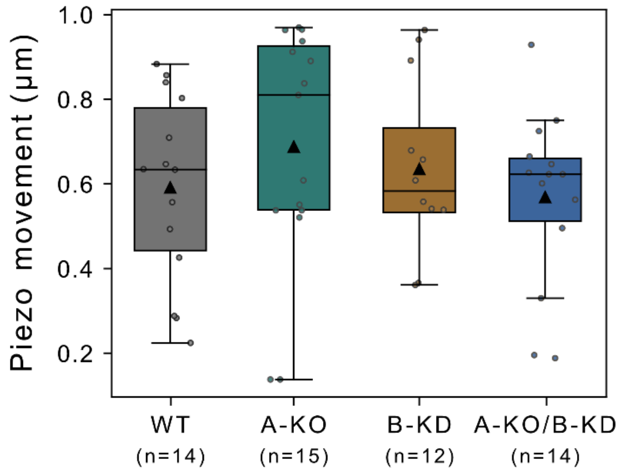


**Fig. S6.** Relative piezo movement from the contact point until reaching 150 nN setpoint for cells with varied lamina compositions: WT, A-KO, B-KO, and A-KO/B-KD. Mean values are shown with triangles in boxplots (N = 3, n ≥ 12 cells). No statistically significant differences were observed between any groups (Kruskal Wallis H-test: H = 1.744, p = 0.627, ns: not significant, Mann-Whitney U tests with Bonferroni correction).

.

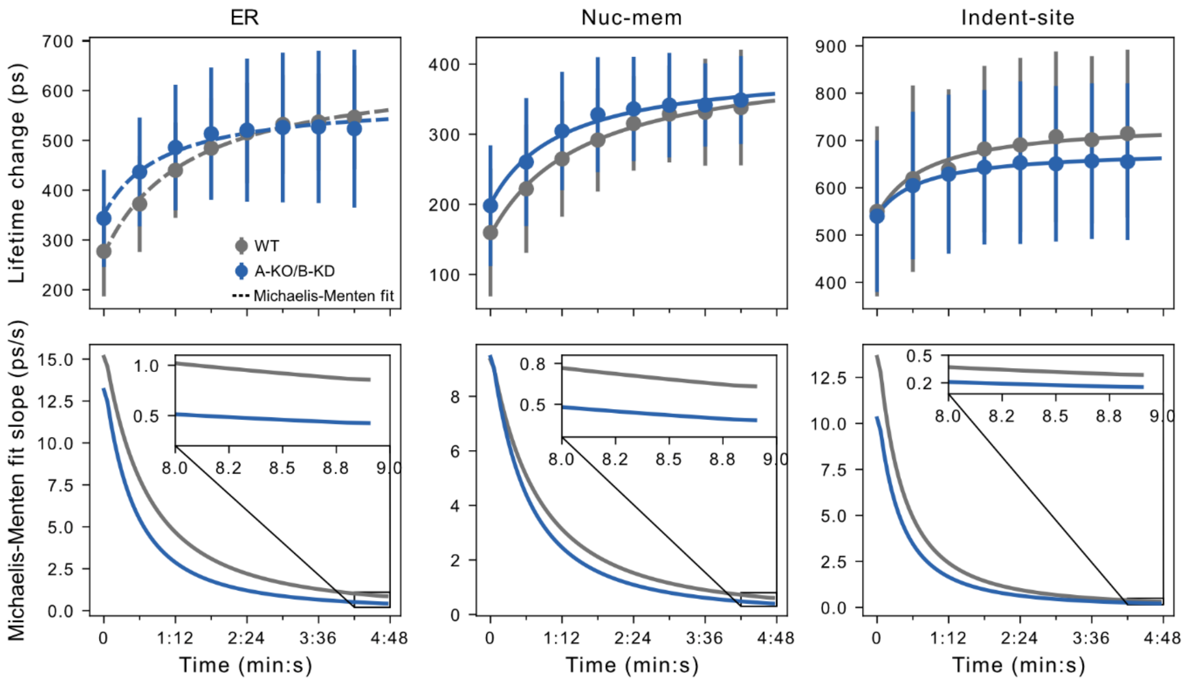


**Fig. S7.** Michaelis-Menten fit to lifetime response (top) and slope value of fitted curve (bottom) in HeLa cells with and without A-type and B-type lamins at the ER, nuclear membrane (Nuc-mem) and indentation site (Indent-site) (Mean ± SD, N = 3, n ≥ 9).

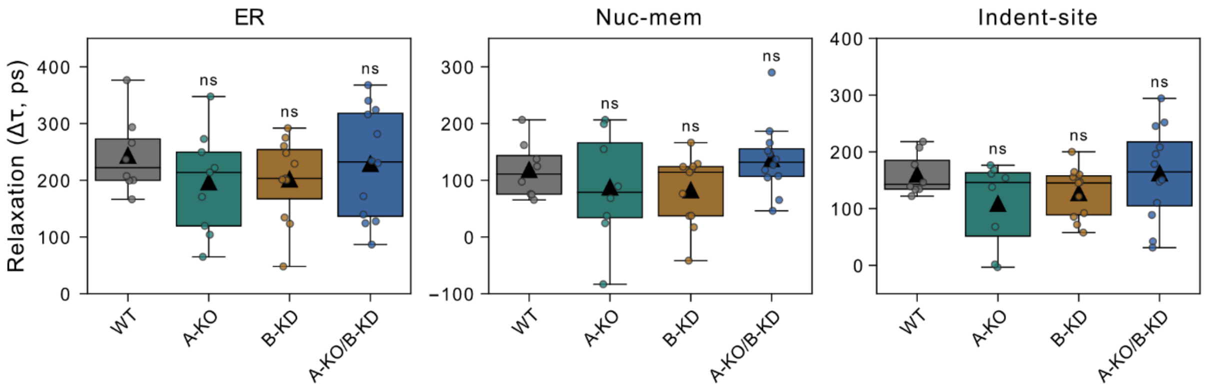


**Fig. S8.** Relaxation behavior of cell with different levels of lamin proteins: WT, A-KO, B-KO and A-KO/B-KD at the ER, nuclear membrane (Nuc-mem) and indentation site (Indent-site) after stimulus. Relaxation is reported as lifetime change 5 min after probe retraction compared to last time point of stimuli. Mean values are shown with triangles in boxplots (N = 3, n ≥ 9, ns: not significant, *t*-test).

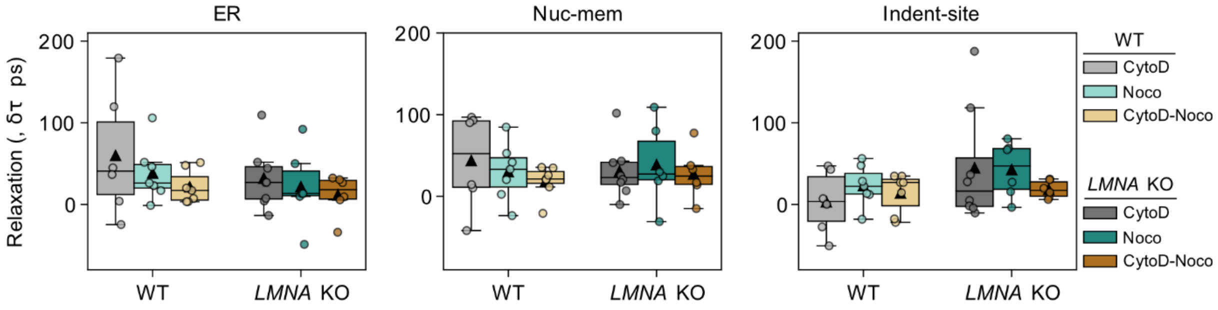


**Fig. S9.** Relaxation behavior of WT and *LMNA* KO cells immediately after mechano-chemical stimuli involving various drugs (CytoD, Noco and CytoD-Noco), at the ER, nuclear membrane (Nuc-mem) and indentation site (Indent-site). Viscoelastic response is composed of lifetime change originating from instantaneous cell response (Elastic) upon indentation and from time-dependent response (Creep) due to AFM probe being in contact (at 150 nN) with cell for around 4 min after different drug injection including CytoD, Noco, CytoD-Noco. Data represent the mean ± SEM (N ≥ 2, n ≥ 6, **p <0.01, *p < 0.05, reported p-values of 0.05 ≤ p < 0.1, t-test with Holm correction). Statistical significance shown on the bars indicates differences between WT and *LMNA* KO cells, emphasizing variations in either the elastic or viscous components under specific drug conditions. The significance levels adjacent to dashed or solid lines signify disparities in the creep or elastic segments of the response, respectively. Relaxation is reported as lifetime change following the retraction of probe compared to last time point of stimuli. Mean values are shown with triangles in boxplots (N ≥ 2, n ≥ 6, no significant differences were captured from *t*-test with Holm correction).
